# Synaptic cell adhesion molecule *Cdh6* identifies a class of sensory neurons with novel functions in colonic motility

**DOI:** 10.1101/2024.08.06.606748

**Authors:** Julieta Gomez-Frittelli, Gabrielle Devienne, Lee Travis, Melinda A. Kyloh, Xin Duan, Tim J. Hibberd, Nick J. Spencer, John R. Huguenard, Julia A. Kaltschmidt

## Abstract

Intrinsic sensory neurons are an essential part of the enteric nervous system (ENS) and play a crucial role in gastrointestinal tract motility and digestion. Neuronal subtypes in the ENS have been distinguished by their electrophysiological properties, morphology, and expression of characteristic markers, notably neurotransmitters and neuropeptides. Here we investigated synaptic cell adhesion molecules as novel cell type markers in the ENS. Our work identifies two Type II classic cadherins, *Cdh6* and *Cdh8,* specific to sensory neurons in the mouse colon. We show that *Cdh6+* neurons demonstrate all other distinguishing classifications of enteric sensory neurons including marker expression of *Calcb* and *Nmu*, Dogiel type II morphology and AH-type electrophysiology and *I_H_* current. Optogenetic activation of *Cdh6+* sensory neurons in distal colon evokes retrograde colonic motor complexes (CMCs), while pharmacologic blockade of rhythmicity-associated current *I_H_* disrupts the spontaneous generation of CMCs. These findings provide the first demonstration of selective activation of a single neurochemical and functional class of enteric neurons, and demonstrate a functional and critical role for sensory neurons in the generation of CMCs.

**One-Sentence Summary:** Intrinsic sensory neurons of the enteric nervous system in the mouse distal colon exclusively express synaptic cell adhesion molecules *Cdh6* and *Cdh8,* evoke retrograde colonic motor complexes (CMCs) when stimulated, and possess rhythmicity-associated *I_H_* current, involved in producing spontaneous CMCs.

## Main Text

### Introduction

Sensory signaling within the gastrointestinal (GI) tract plays a critical role in the autonomous regulation of digestion. The GI tract is the only internal organ system containing its own sensory neurons. Intrinsic primary afferent neurons (IPANs) detect relevant stimuli through chemo- and mechano-sensation and direct appropriate GI functions via downstream components of the enteric nervous system (ENS), including ascending and descending interneurons, and excitatory and inhibitory motor neurons (*1*). These neuronal subtypes have begun to be distinguished morphologically, electrophysiologically, and by marker expression, classically especially of neurotransmitters (*2*, *3*), and together this information has provided an opening to characterize individual neuron subtype function within the GI tract.

Synaptic cell adhesion molecules define neuronal subtype connectivity within many regions of the CNS. Type II Cadherins are a family of synaptic cell adhesion molecules with combinatorial expression in multiple neural circuits of the CNS, including retina, limbic, olivonuclear, and auditory projection systems (*4–6*). Type II Cadherins bind homophilically by expression of the same cadherin at both the pre- and post-synapse, which stabilizes developing synapses between correct partners while incorrect synapses are pruned away (*7*, *8*). Recent RNA-Seq studies of human and mouse ENS have identified synaptic cell adhesion molecules, including Type II Cadherins, expressed in enteric neuronal subtypes (*9–11*). However, the specificity of their expression has yet to be harnessed to assess neuronal subtype-specific function in the ENS.

Here we identify Type II Cadherin, *Cdh6*, as a novel marker for IPANs of the colonic ENS. We demonstrate the sensory identity of *Cdh6* neurons by immunohistochemical, morphological and neurophysiological classification. *Cdh6* neurons express IPAN markers *Calcb* and *Nmu*. Sparse labeling of individual IPANs reveals they project mainly circumferentially and branch extensively in myenteric ganglia. Whole cell patch clamp recordings of sensory neurons *in situ* reveal action potential (AP) slow after-hyperpolarizations characteristic of IPANs, and hyperpolarization-activated cationic current (*I_H_*), a rhythmicity indicator in thalamocortical and other systems (*12*). Using a *Cdh6* genetic mouse model, we show that optogenetic activation of distal colon IPANs is sufficient to evoke retrograde CMCs, while pharmacologic block of *I_H_* in IPANs disrupts colonic rhythmicity and reversibly abolishes spontaneous CMCs.

### Results

#### Expression of the type II classic cadherin *Cdh6* in colonic IPANs

To identify Cadherins expressed in enteric neuronal subtypes in mouse, we screened recently published RNA-Seq data (*10*, *11*) for Classic Type II cadherin expression. *Cdh6* and *Cdh8* appeared to be restricted to IPAN subsets in both small intestine and colon (*10*, *11*). *Cdh9* was previously identified in a separate population of IPANs in the small intestine (*9–11*). We validated *Cdh6* and *Cdh8* expression by RNAscope *in situ* hybridization in the myenteric plexus, which contains the enteric motility circuitry. *Cdh6* mRNA was expressed in 14.7 ± 0.8% of myenteric neurons in small intestine (jejunum) and in 6.8 ± 0.3% of myenteric neurons in distal colon (mean ± SEM) [Fig.1A-C]. *Cdh8* was almost exclusively co-expressed in *Cdh6+* neurons, although at a much lower level of detection [Fig.1D-H]. We therefore focused our further analysis on *Cdh6+* neurons.

**Fig. 1.**
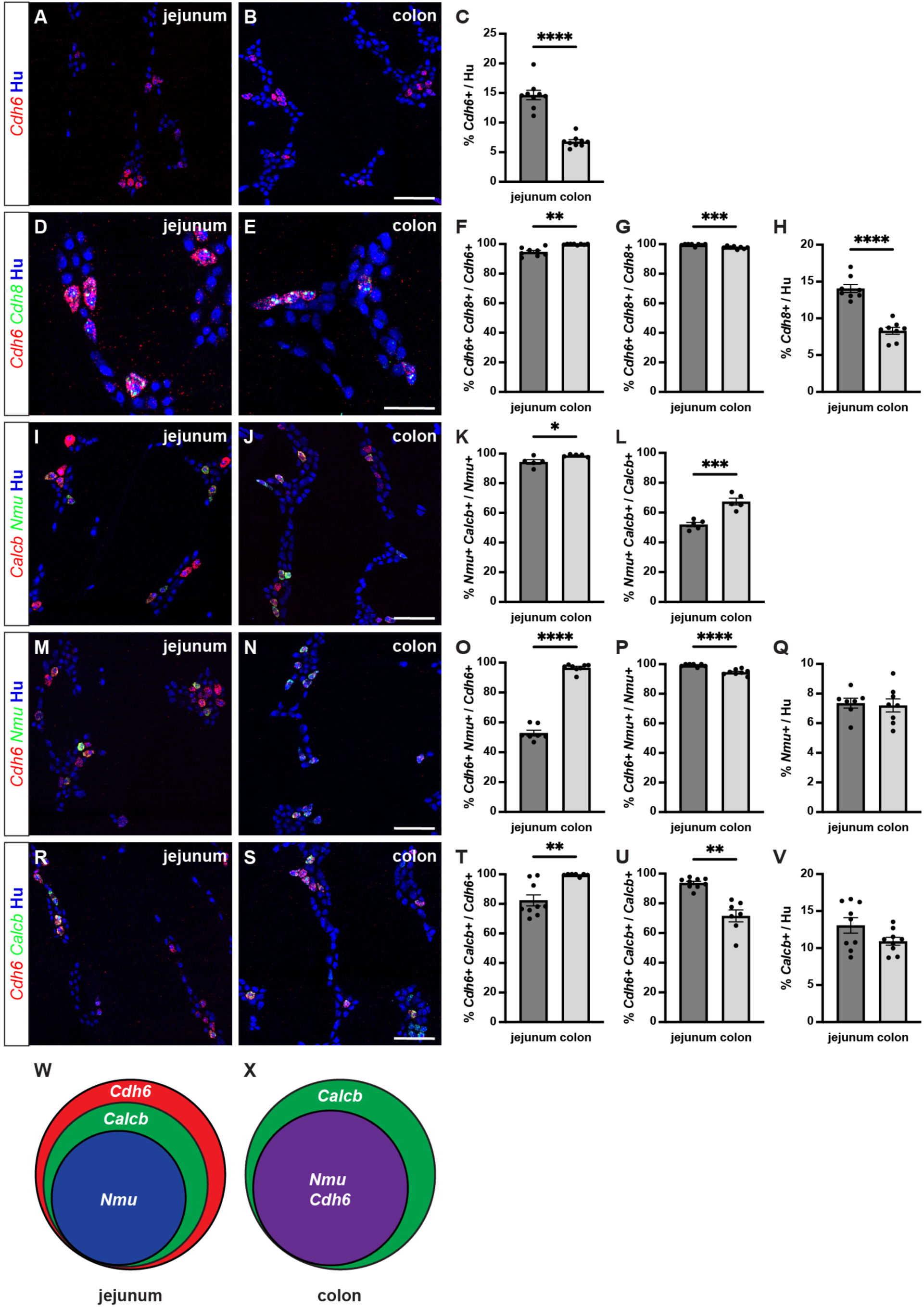
*Cdh6* expression overlaps with IPAN markers *Calcb* and *Nmu*. (**A, B**) Representative images of jejunum (A) and distal colon (B) myenteric plexus labeled with HuC/D (IHC) (blue) and *Cdh6* (RNA) (red). (**C**) Proportion of total HuC/D neurons positive for *Cdh6* (jejunum, n = 9; distal colon, n = 9). (**D, E**) As in (A, B) for HuC/D (IHC) (blue), *Cdh6* (RNA) (red), and *Cdh8* (RNA) (green). (**F**) Proportion of *Cdh6+* neurons positive for *Cdh8* (jejunum, n = 8; distal colon, n = 8). (**G**) Proportion of *Cdh8+* neurons positive for *Cdh6* (jejunum, n = 8; distal colon, n = 8). (**H**) Proportion of total HuC/D neurons positive for *Cdh8* (jejunum, n = 8; distal colon, n = 8). (**I, J**) As in (A, B) for HuC/D (IHC) (blue), *Calcb* (RNA) (red), and *Nmu* (RNA) (green). (**K**) Proportion of *Nmu+* neurons positive for *Calcb* (jejunum, n =5 ; distal colon, n = 5). (**L**) Proportion of *Calcb+* neurons positive for *Nmu* (jejunum, n = 5; distal colon, n = 5). (**M, N**) As in (A, B) for HuC/D (IHC) (blue), *Cdh6* (RNA) (red), and *Nmu* (RNA) (green). (**O**) Proportion of *Cdh6+* neurons positive for *Nmu* (jejunum, n = 7; distal colon, n = 8). (**P**) Proportion of *Nmu+* neurons positive for *Cdh6* (jejunum, n = 7; distal colon, n = 8). (**Q**) Proportion of total HuC/D neurons positive for *Nmu* (jejunum, n = 7; distal colon, n = 8). (**R, S**) As in (A, B) for HuC/D (IHC) (blue), *Cdh6* (RNA) (red), and *Calcb* (RNA) (green). (**T**) Proportion of *Cdh6+* neurons positive for *Calcb* (jejunum, n = 9; distal colon, n = 7). (**U**) Proportion of *Calcb+* neurons positive for *Cdh6* (jejunum, n = 9; distal colon, n = 7). (**V**) Proportion of total HuC/D neurons positive for *Calcb* (jejunum, n = 9; distal colon, n = 9). (**W, X**) Schematic of marker overlap in jejunum (W) and distal colon (X). Scale bar represents 100 μm for (A, B, I, J, M, N, R, S), 50 μm for (D, E). All charts (mean ± SEM). * p<0.05; ** p<0.01; *** p<0.001; **** p<0.0001.

To confirm IPAN identity of *Cdh6+* neurons, we first established the differential expression of two putative and broadly used markers of IPANs, *Calcb* and *Nmu* (*3*, *9*, *11*). We found that all *Nmu+* neurons co-express *Calcb* in both jejunum and distal colon [Fig.1I-K]. In contrast, only about half of *Calcb+* neurons in the jejunum and two-thirds in the distal colon co-express *Nmu* [Fig.1L].

We next assessed co-expression of *Cdh6* with *Calcb* and *Nmu.* In the jejunum, we found that nearly all *Nmu+* neurons and *Calcb+* neurons express *Cdh6* [Fig.1P, U], though only about three-quarters of all *Cdh6+* neurons express *Calcb* and only about half express *Nmu* [Fig.1O, T]. In contrast, in the distal colon, while *Cdh6* is only expressed in about two-thirds of all *Calcb+* neurons [Fig.1U], nearly all *Cdh6+* neurons express *Nmu* and *Calcb* [Fig.1O, T]. Taken together, our data show that in the myenteric plexus, *Cdh6* is expressed exclusively in *Calcb+/Nmu+* IPANs in the mouse distal colon [Fig.1W, X].

#### Mouse colonic IPANs display AH-type electrophysiology and *I_H_* current

We next assessed the electrophysiological properties of *Cdh6+* IPANs. We focused our analysis on the colon, and for ease of neuron tracing, took advantage of a genetic strategy to sparsely label *Cdh6+* neurons. Previous studies of Hb9:GFP transgenic mice have shown that due to the inserted transgene’s proximity to *Cdh6*, *Cdh6+* neurons can express eGFP (*13*). Hb9:GFP+ neurons were rare and projected extensively throughout the myenteric plexus [Fig.2A-C]. *In situ* hybridization confirmed *eGFP* expression was limited to a small fraction (3.5 ± 0.8%) of *Cdh6+* colonic neurons [Fig.2D, F-H], and all *eGFP+* neurons expressed *Cdh6* [Fig.2E].

**Fig. 2.**
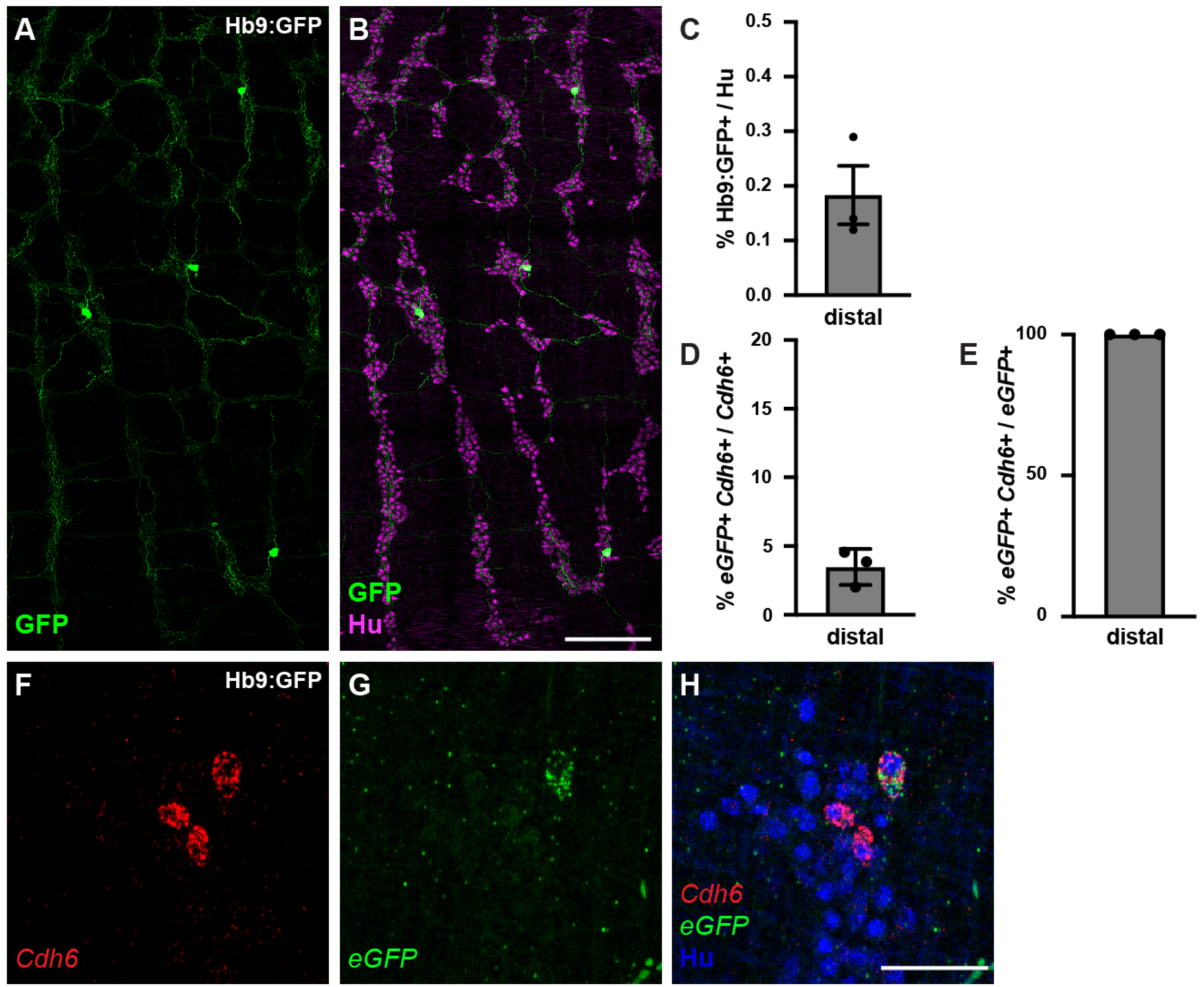
Hb9:GFP+ is expressed in a small proportion of *Cdh6+* colon myenteric neurons. (**A, B**) Representative images of Hb9:GFP+ distal colon myenteric plexus labeled with HuC/D (IHC) (magenta) and GFP (green). (**C**) Proportion of total distal colon HuC/D neurons positive for GFP (n = 3). (**D**) Proportion of distal colon *Cdh6+* neurons positive for *eGFP* (n = 3). (**E**) Proportion of distal colon *eGFP+* neurons positive for *Cdh6* (n=3). (**F-H**) Representative images of Hb9:GFP+ distal colon myenteric plexus labeled with HuC/D (IHC) (blue), *Cdh6* (RNA) (red), and e*GFP* (RNA) (green). Scale bar represents 200 μm for (A, B), 50 μm for (E-G). All charts (mean ± SEM).

We developed a protocol to perform whole-cell patch-clamp recordings (*14*) in Hb9:GFP+ colonic neurons in the distal colon. Membrane capacitance reflecting overall size of these cells was 32 ± 8.7 pF [Fig.3D]; their resting membrane potential (RMP) was −49.4 ± 2.9 mV [Fig.3C]. The input resistance (R_in_) was 393 ± 54.7 MΩ [Fig.3E] as computed from the slope of the voltage-current (V-I) relationship. All patched neurons had large-amplitude action potentials (AP; 72 ± 2.5 mV, [Fig.3F, E]) with threshold of −26.4 ± 0.9 mV [Fig.3L] and a half width of 1.2 ± 0.1 ms [Fig.3H] elicited at rheobase (20 ± 4.5 pA, [Fig.3G]), each followed by an afterhyperpolarization (AHP= −67.7 ± 0.9 mV, [Fig.3K]). In addition, the first derivative of membrane voltage during the action potential (dV/dt, [Fig.3J]) exhibited an inflection during the repolarization phase, suggesting the presence of fast and slow AP repolarization mechanisms [Fig.3F]. Patched Hb9:GFP+ neurons generally could not sustain repetitive APs in response to long depolarizing current pulses. All patched neurons responded to hyperpolarizing current pulses with a time-dependent membrane potential sag (−6.3 ± 1.2 mV, [Fig.3B, M]), and a rebound depolarization following the hyperpolarization [Fig.3B, N].

**Fig. 3.**
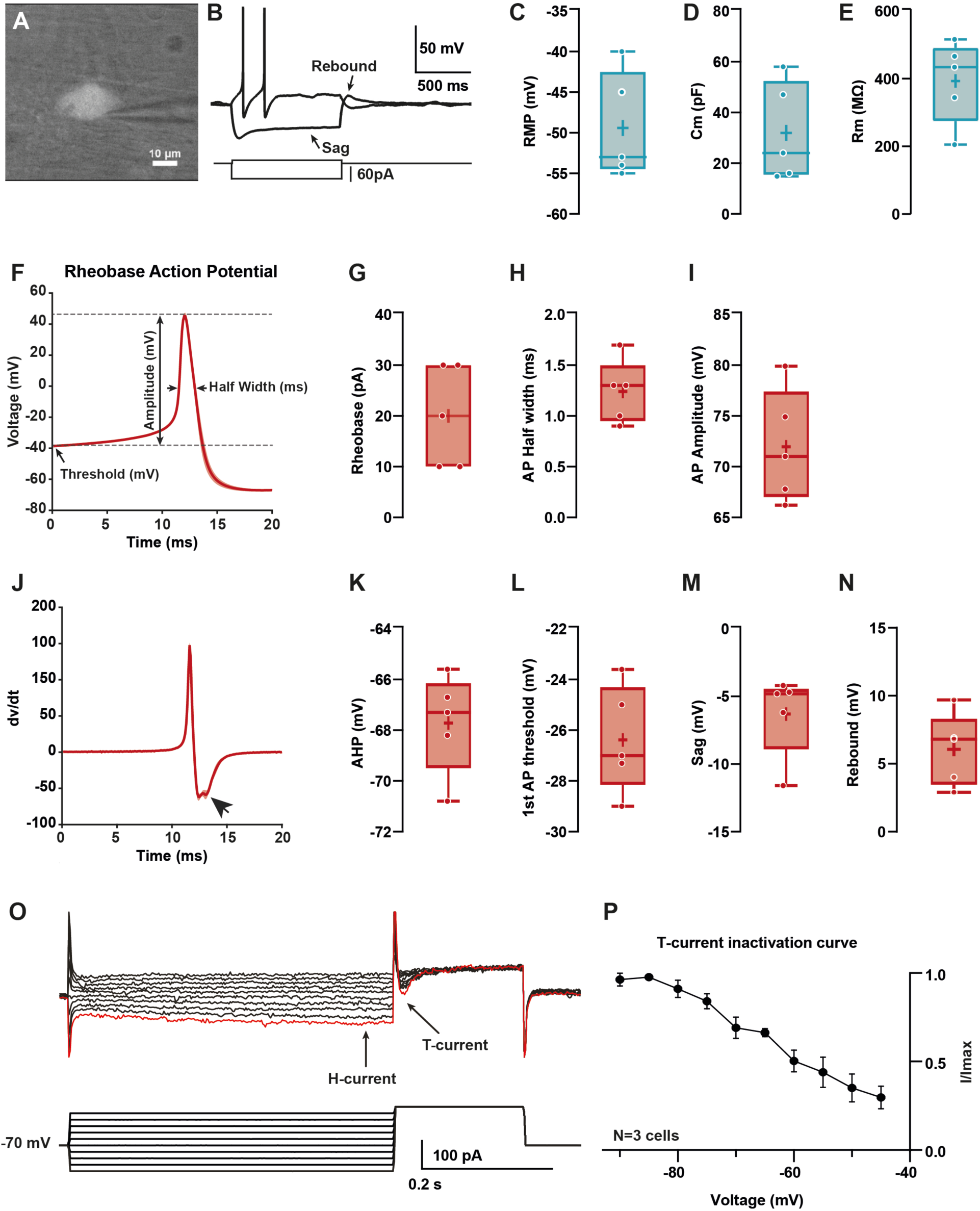
Hb9:GFP+ distal colon neurons have AH electrophysiological characteristics. (**A**) IR videomicroscopy image of an Hb9:GFP distal colon neuron that presented a large soma located in a ganglion (scale bar, 10 μm). (**B**) Current-clamp recordings of the same neuron in (A) obtained in response to application of current pulse (bottom traces) of −50 pA, and + 10 pA. Note the presence of a sag and a post-hyperpolarization rebound depolarization. (**C-E**) Box-and-whisker plots of cellular properties of recorded neurons. (C) resting membrane potential (RMP), (D) capacitance (Cm), and (E) membrane resistance (Rm) (N = 5). (**F**) Averaged traces of the first spike (rheobase action potential) after a depolarization step of 1s. (**J**) Averaged derivative traces of the first spike (rheobase action potential). An inflection on the repolarizing phase is observed in the first derivative (arrow). (**G-I, K-N**) Box-and-whisker plots of electrophysiological properties of recorded neurons; rheobase action potential (AP, G-I) (G) current threshold, (H) half-width, (I) amplitude, (K) afterhyperpolarization (AHP), and (L) threshold. (M, N) non-AP properties sag (mV) and rebound (mV). (**O**) H and T currents in recorded neurons. Top: example of currents obtained from voltage protocol. Bottom: 500 ms hyperpolarizations ranging from −90 to −45 for 500 ms followed by depolarizing to −40 mV. Hyperpolarizations evoked slowly activating inward current (H-Current, arrow), followed by a transient inward current upon post-conditioning step to −30mV (T- current, arrow) Largest T and H currents were obtained with the most hyperpolarized potentials (red trace). (**P**) Normalized peak I_T_ plotted versus holding potential to obtain the I/Imax curve (N = 3/3). Scale bar represents 10 μm for (A).

Consistent with these findings of sag and rebound, voltage clamp step hyperpolarizations revealed *I_H_* (hyperpolarization-activated current, [Fig 3.N]) and, upon repolarization, *I_T_* (transient inward presumed Ca^2+^ current) (*15*) in terms of its kinetics [Fig 3.O] and steady-state inactivation [Fig.3O, P]. Taken together, these results show that Hb9:GFP+ / *Cdh6+* distal colonic neurons have AP after-hyperpolarizing (AH)-type electrophysiology typical of IPANs, including rhythm generating currents *I_H_* and *I_T_* (*16–18*).

#### Colonic IPANs have Dogiel type II morphology and abundant projections throughout the myenteric plexus

To visualize the morphology and projections of individual patched Hb9:GFP+ neurons, we included biocytin in the internal solution for post-fixation single-cell tracing [Fig.4]. Patched Hb9:GFP+ neurons displayed Dogiel type II morphology (*19*), with large smooth cell somas and multiple branching neurites. Projections were mainly circumferential and extensively branched within myenteric ganglia. Thus, Hb9:GFP+ neurons display morphological features characteristic of IPANs (*2*, *19*).

**Fig. 4.**
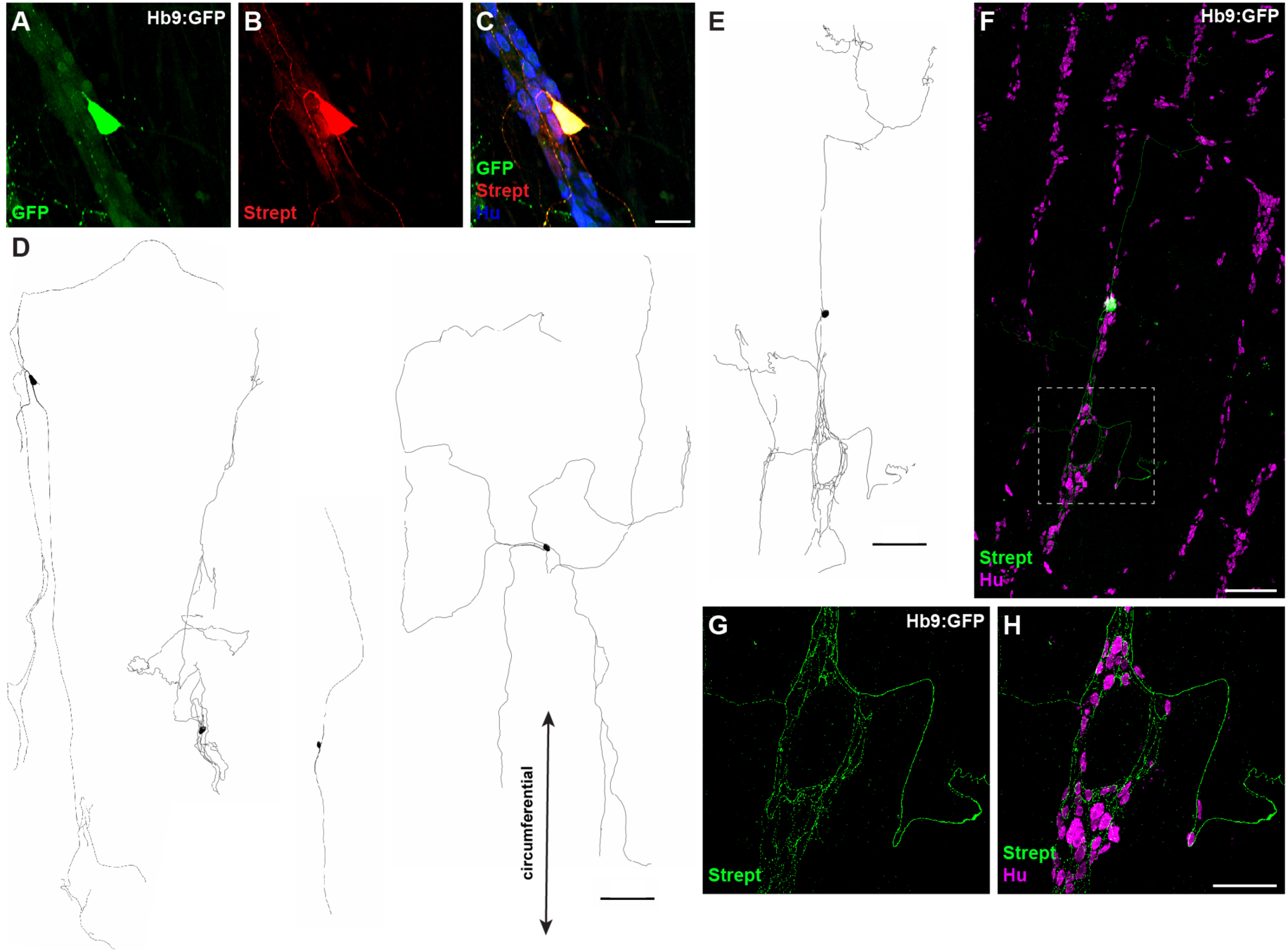
Hb9:GFP+ distal colon neurons have circumferential branching projections. (**A-C**) Representative images of Hb9:GFP+ distal colon myenteric plexus labeled with HuC/D (blue), streptavidin (red), and GFP (green). (**D, E**) Tracings of Hb9:GFP+ distal colon neurons filled with biocytin during whole cell patch clamp recording. (**F**) Image of patched and filled Hb9:GFP+ distal colon neuron traced in (E). (**G, H**) Inset of (F). Scale bar represents 40 μm for (A-C), 200 μm for (D-F), 100 μm for (G, H).

To further visualize the full extent of IPAN circuitry in the myenteric plexus, we intercrossed Cdh6^CreER^ (*20*) and LSL-tdTomato (Ai14) (*21*) mice and induced Cre expression at 5-8 weeks of age [Fig.5A, B]. We confirmed tdTomato+ labeling in myenteric *Cdh6+* neurons [Fig.5E, F], representing about 5% of the total neuronal population [Fig.5G]. tdTomato+ neurons had large cell somas (major axis, 27.8 ± 0.7 µm; minor axis, 15.9 ± 0.4 µm) [Fig.5D]. All ganglia of the myenteric plexus were densely innervated by tdTomato+ fibers [Fig.5A, B], which also projected into the circular muscle. We noted additional tdTomato+ labeling of some putative longitudinal and circular muscle cells [Fig.5A, B]. Taken together, our data reveal an IPAN array that spans the entire motility circuitry of the colonic myenteric plexus.

**Fig. 5.**
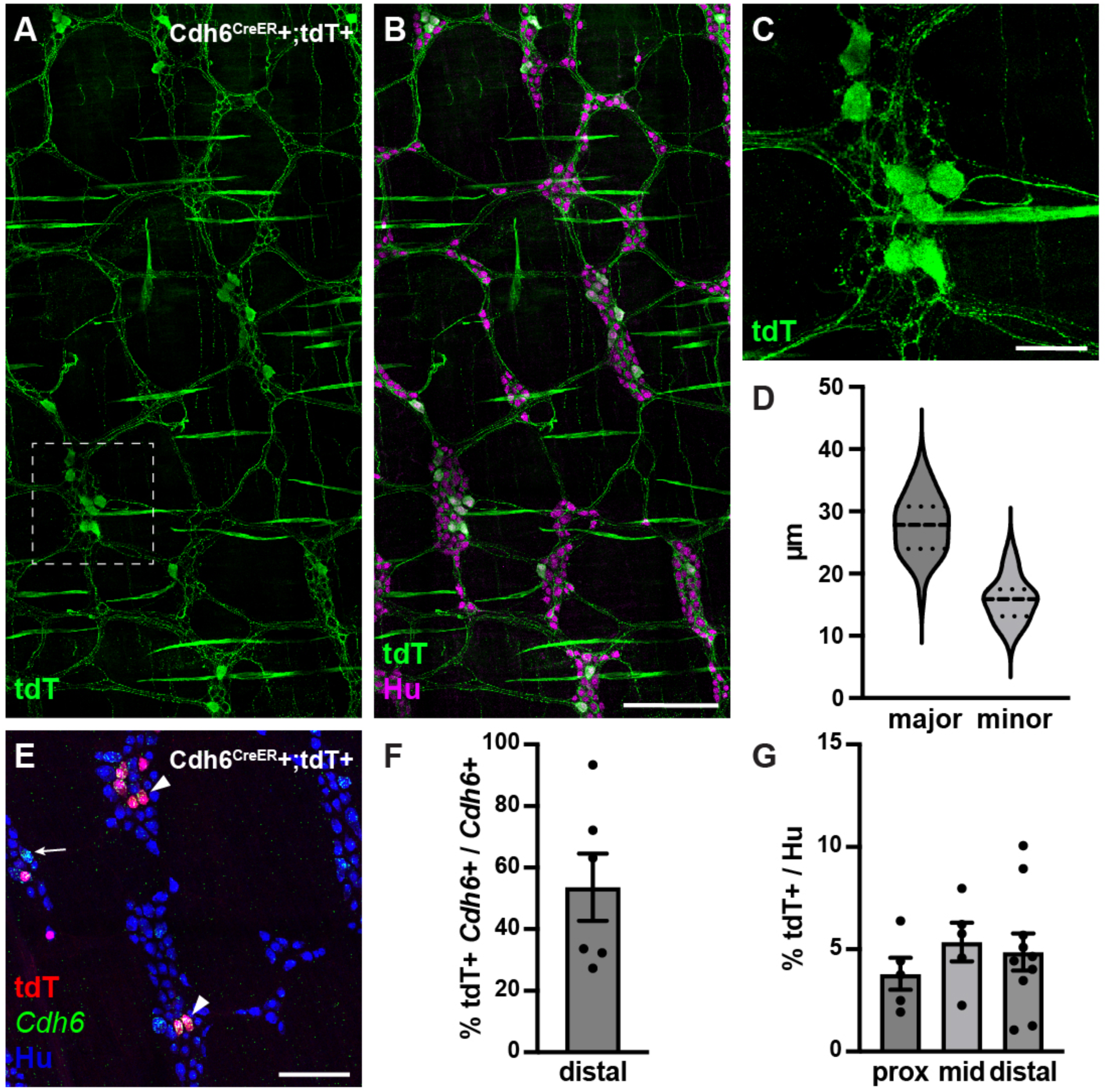
Cdh6^CreER^+/ tdTomato*+* neurons have Dogiel type II morphology. (**A, B**) Representative images of Cdh6^CreER^+;tdTomato+ distal colon myenteric plexus labeled with HuC/D (IHC) (magenta) and tdTomato (IHC) (green). (**C**) Inset of (A). (**D**) Dimensions of tdTomato+ neurons (major and minor axes) (N = 73; n = 3). (**E**) Representative image of Cdh6^CreER^+;tdTomato+ distal colon myenteric plexus labeled with HuC/D (IHC) (blue), tdTomato (IHC) (red), and *Cdh6* (RNA) (green). Arrowheads indicate *Cdh6*+ / tdTomato+ cells; arrowhead, *Cdh6*+ / tdTomato-negative cell. (**F**) Proportion of *Cdh6+* distal colon neurons positive for tdTomato (n = 6). (**G**) Proportion of total HuC/D neurons positive for tdTomato (proximal colon, n = 5; mid colon, n = 5; distal colon, n = 10). Scale bar represents 100 μm for all images. All charts (mean ± SEM).

#### Optogenetic activation of distal colon IPANs evokes CMCs

In thalamocortical relay neurons, *I_H_* contributes to intrinsic slow rhythmic burst firing at 1-2Hz (*22*), but the function of *I_H_* in colonic IPANs is not known. During colonic motor complexes (CMCs), large regions of the ENS oscillate in synchrony at 1-2Hz to generate traveling contractions along the colon (*23*). Recent calcium imaging studies have shown that IPANs participate, along with all other subtypes of enteric neurons, in this synchronized oscillatory firing (*24*). Furthermore, our electrophysiological studies confirm the presence of *I_H_* in mouse colonic IPANs [Fig.3N]. However, the role of IPANs in spontaneous CMCs is not well understood.

To interrogate the functional role of *Cdh6+* IPANs in CMCs, we performed *ex vivo* colonic contraction force recordings in conjunction with optogenetic activation. To express ChR2-eYFP in *Cdh6+* cells, we intercrossed Cdh6^CreER^ and LSL-ChR2-eYFP (Ai32) (*25*) mice and induced Cre expression at 5-8 weeks of age [Fig.6A-C]. In Cdh6^CreER^+;ChR2-eYFP+ colon preparations we observed spontaneous CMCs at regular intervals of about 3-5 minutes. Blue light stimulation of *Cdh6+* IPANs in distal colon 60-90 seconds after a spontaneous CMC (“control” CMC) resulted in an evoked, premature CMC that began during stimulation (N = 17 stimulations; n = 5 mice)[Fig.6D, E]. Evoked CMCs traveled retrogradely from the distal to the proximal colon. They were similar to spontaneous CMCs in peak amplitude, area under the curve, and duration, though the contractile force (peak amplitude and AUC) was slightly weaker in the proximal colon [Fig.6F-H]. Blue light stimulation in proximal or mid colon failed to generate CMCs (n = 5/5, data not shown). In comparison, stimulation in control Cdh6^CreER^- negative;ChR2-eYFP+ colons failed to evoke any CMCs (N = 28 stimulations; n = 7 mice)[Fig.S1].

**Fig. 6.**
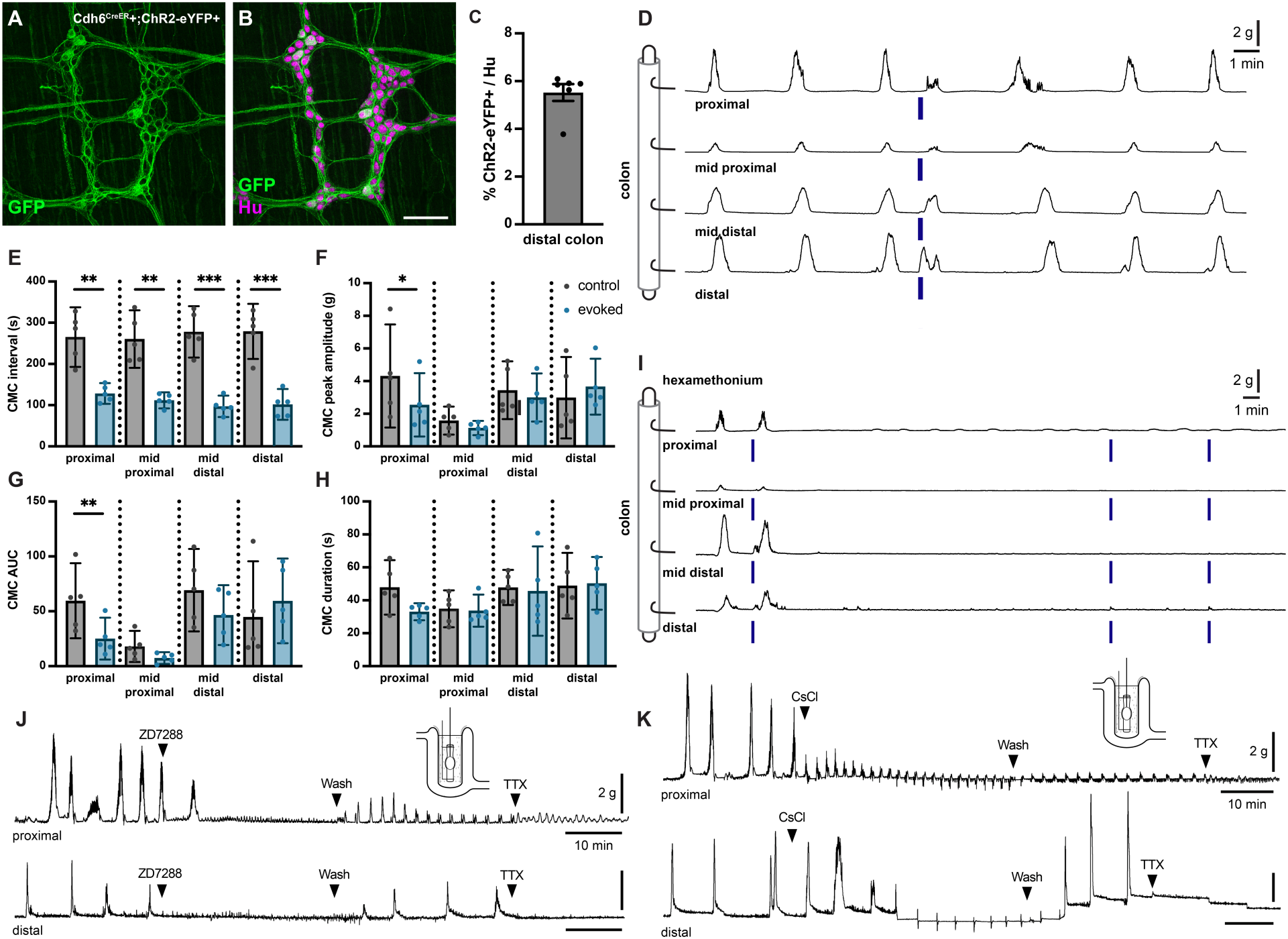
Optogenetic stimulation of distal colonic *Cdh6+* neurons evokes CMCs, while pharmacologic blockade of *I_H_* abolishes spontaneous CMCs. (**A, B**) Representative images of Cdh6^CreER^+;ChR2-eYFP+ distal colon myenteric plexus labeled with HuC/D (magenta) and GFP (green). (**C**) Proportion of total distal colon HuC/D neurons positive for ChR2-eYFP (n = 6). (**D**) Representative force traces. Blue bars indicate timing of light stimulation. LEDs placed distal to distal hook. (**E**) CMC intervals recorded from force traces. Evoked (blue) intervals represent the time from the prior spontaneous CMC before stimulation to the evoked CMC following stimulation. Control (grey) intervals represent the time between the spontaneous CMC prior to stimulation and the previous spontaneous CMC (n = 5). Paired t test, one tailed. (**F**) CMC peak amplitude recorded from force traces. Evoked (blue) indicates the evoked CMC following stimulation. Grey (control) indicates the spontaneous CMC prior to stimulation (n = 5). Paired t test, two tailed. (**G**) CMC AUC (area under the curve). Evoked (blue) and control (grey) as in (F) (n = 5). Paired t test, two tailed. (**H**) CMC duration. Evoked (blue) and control (grey) as in (F) (n = 5). Paired t test, two tailed. (**I**) Representative force traces. Hex indicates addition of 300 µM hexamethonium. Blue bars indicate timing of light stimulation. LEDs placed distal to distal hook (n = 5/5). (**J**) Representative force traces on tethered pellets. First arrowhead indicates addition of 10 µM ZD7288. Second arrowhead indicates washout in Krebs. Third arrowhead indicates addition of 1 µM TTX. ZD7288 abolished CMCs in both proximal and distal colon (n = 6/6, p = 0.0022, Fisher’s exact test). Washout in Krebs restored CMCs in both proximal and distal colon (n = 6/6, p = 0.0022, Fisher’s exact test). (**K**) As in (J). First arrowhead indicates addition of 2 mM CsCl. Second arrowhead indicates washout in Krebs. Third arrowhead indicates addition of 1 µM TTX. Typical CMC production was impaired or altered by CsCl (proximal colon, n = 5/6, p = 0.0152; distal colon, n = 6/6, p = 0.0022, Fisher’s exact test): increased frequency (proximal colon, n = 5/6, p = 0.0152; distal colon, n = 6/6, p = 0.0022, Fisher’s exact test), decreased in amplitude (proximal colon, n = 5/6, p = 0.0152; distal colon, n = 4/6, p = 0.0606, Fisher’s exact test); retrograde force (proximal colon, n = 2/6; distal colon, n = 2/6). Scale bar represents 100 μm for (A, B). * p<0.05; ** p<0.01; *** p<0.001.

CMCs have previously been shown to depend on nicotinic cholinergic transmission (*24*). We performed optogenetic stimulation in the presence of hexamethonium, a blocker of nicotinic cholinergic transmission. Spontaneous CMCs were abolished in hexamethonium, and CMCs could not be evoked by optogenetic stimulation [Fig.6I]. We conclude that activation of *Cdh6+* distal colon IPANs evokes retrograde-traveling but otherwise characteristic and hexamethonium-sensitive CMCs.

#### Blockade of *I_H_* current in colonic IPANs disrupts CMC production

To determine whether *I_H_* in IPANs may contribute to oscillatory firing driving CMCs, we measured colonic contraction force on a tethered pellet in the presence of *I_H_* blockers ZD7288 or CsCl (*26*, *27*). ZD7288 blocks *I_H_* in all IPANs, including *Cdh6*+ IPANs. Spontaneous CMCs were recorded in all preparations of both proximal and distal colon prior to drug application [Fig.7]. Addition of 10 µM ZD7288 to the recording chamber abolished spontaneous CMCs and washout of ZD7288 recovered spontaneous CMC activity [Fig.6J, Fig.S2]. Addition of 2mM CsCl also impaired or altered spontaneous rhythmic production of typical CMCs [Fig.6K, Fig.S3]; rhythmic contractions increased in frequency (proximal colon, n = 5/6; distal colon, 6/6), decreased in amplitude (proximal colon, n = 5/6; distal colon, n = 4/6, p = 0.0606), or in some cases even included significant retrograde force components (proximal colon, n = 2/6; distal colon, n = 2/6). We conclude that pharmacologic blockade of *I_H_* in IPANs impairs the production of CMCs in the mouse colon.

### Discussion

Our study shows that in the myenteric plexus, *Cdh6* is expressed exclusively in *Calcb+/Nmu+* IPANs located in the mouse distal colon, while in the small intestine, *Cdh6* is also expressed in some *Calcb+/Nmu-* and *Calcb-/Nmu-* neurons. We confirm the IPAN identity of *Cdh6+* distal colonic neurons by electrophysiological recordings revealing AH-type signature, and single neuron tracings showing Dogiel type II morphology. Finally, we demonstrate that activation of IPANs in the distal colon evokes retrograde CMCs, while pharmacologic blockade of *I_H_*, a rhythmicity associated current we show also present in mouse colonic IPANs, disrupts spontaneous CMC generation.

Together with *Cdh8*, which we show to be co-expressed with *Cdh6,* our study validates two new adhesion molecules specific to IPANs in the distal colon. Notably, these markers show a different expression pattern than another cadherin, *Cdh9*, which is exclusively expressed in the small intestine in mouse, not in the colon (*9*). *Cdh9* is expressed in a subset of IPANs not expressing either *Calcb* or *Nmu* (*3*, *9*, *11*). It is possible that in the small intestine *Cdh6* and *Cdh9* mark some of the same neurons. However, RNA-Seq data from two separate studies suggest this is unlikely (*10*, *11*). We conclude that *Cdh6/Cdh8* IPANs are a separate population from *Cdh9* IPANs. Finally, our study was limited to the myenteric plexus, containing motility circuitry. IPANs are also present in the submucosal plexus, and it would be interesting to investigate *Cdh6* expression and neuronal subtype in the SMP.

Though IPANs are positioned to initiate motility by activating other subtypes in enteric circuitry, their role in spontaneous CMCs has been debated, as CMCs can occur in the absence of luminal contents, without any apparent stimulus for IPANs to sense, or upon stimulation of nitrergic populations (*2*, *28*). Only recently has evidence emerged to suggest that IPANs participate in oscillatory rhythmic firing of the ENS during CMCs, and that activation of neuronal subtypes expressing calretinin, including IPANs, together can evoke CMCs (*24*, *29*). Our results demonstrate that excitement of IPANs alone in the distal colon is capable of producing retrograde CMCs. The exact mechanism causing CMC generation at an established and controlled frequency of once every few minutes, without evident stimulus, remains to be determined.

A major observation was that optogenetic stimulation of *Cdh6*+ neurons readily evoked CMCs from the distal colon, but never from the proximal colon. Failure of proximal colonic stimulation to evoke CMCs may reflect inhibition by IPAN recruitment of descending pathways; or conversely, high efficacy of stimulation from distal colon may reflect bias toward activation of ascending excitatory cholinergic pathways. Our classification of *Cdh6+* neurons as IPANs in the distal colon may not extend to the proximal colon; thus, we cannot extrapolate that *Cdh6*+ optogenetic activation in the colon is restricted to IPANs as in the distal colon. In addition, our stimulus paradigm in the proximal colon may not have activated enough neurons based on light density and illumination. In previous studies, optogenetic stimulation of calretinin-expressing neurons or choline acetyl transferase (ChAT)-expressing neurons (both of which include IPANs, interneurons and motor neurons) elicited anterograde CMCs regardless of stimulus location, including proximal colon (*29*, *30*). Proximal colon stimulation of nitric oxide synthase-expressing neurons also elicited CMCs (*28*). These studies, as noted, each activated multiple neuronal classes employing the same neurotransmitter. How broad and non-specific activation of such disparate neuron classes readily evokes CMCs remains unclear.

Spontaneous and evoked CMCs were abolished in hexamethonium, confirming that CMC synchronous firing is dependent on nicotinic cholinergic transmission (*24*). This reinforces that nicotinic cholinergic transmission is required for greater activation and synchrony of the entire ENS motility network to generate CMCs.

Through our electrophysiological investigation of IPANs in the distal colon using voltage clamp, our work reveals the presence of two voltage gated ion conductances and their underlying currents, *I_T_* and *I_H_*. Slow AHP, *I_H_* and *I_T_* have been previously identified as distinguishing characteristics of IPANs in rat and guinea pig (*16–18*). Although prior studies noted *I_H_* and proposed *I_T_* in guinea pig Dogiel type II neurons (*17*), they have not previously been reported in studies of intact mouse distal colon myenteric plexus due to the conventional reliance on sharp electrode recordings in which voltage clamp is not possible (*31*). The presence of *I_H_* and *I_T_* in thalamocortical relay neurons and other cell types supports intrinsic rhythmicity (*22*, *32*). It is possible that these two currents may similarly promote autonomous rhythmic activity in colonic IPANs.

*I_H_* is conducted through hyperpolarization-activated cyclic nucleotide gated (HCN) channels (*33*). HCN channel family members HCN1 and HCN2 have been shown to be present in mouse distal colonic Dogiel type II neurons (*16*), and RNA-Seq ENS screens similarly indicate their expression in IPANs (*10*, *11*). Knockout of HCN2 in mouse leads to a severe growth restriction phenotype due to malnutrition and GI dysmotility (*34*). Here we demonstrate that pharmacologic blockade of *I_H_*, which we show to be present in IPANs, with two distinct HCN channel blockers ZD7288 and CsCl abolishes CMCs, an otherwise persistent and ongoing pattern of motor activity in the mouse colon. We speculate that *I_H_* in colonic IPANs, as in thalamocortical neurons, plays a role in promoting either rhythmic oscillatory single neuron activity, or network burst firing, or both (*22*, *32*). Blocking *I_H_* may impair individual IPANs’ ability to fire rhythmically, or the ability of IPANs to synchronize into a network burst firing mode. Failure of IPANs to fully activate and synchronize could prevent generation of both (1) synchronized rhythmic myenteric network activation of motility circuits, and (2) the resulting synchronized contractions that sum to much larger contractile forces during CMCs.

Type II cadherins are most commonly homophilic synaptic cell adhesion molecules (*7*, *8*). *Cdh6* can also form heterodimers with *Cdh7, Cdh10,* and *Cdh14* (*35*). However, *Cdh10* and *Cdh14* are very lowly expressed in the colon by single cell RNA sequencing (*10*), and we performed RNAscope for *Cdh7* and did not observe any expression (data not shown). Restricted expression of two Type II cadherins, *Cdh6* and *Cdh8*, to mouse colonic IPANs raises the possibility of these cadherins supporting IPAN-IPAN synaptic connections. While broadly speaking, IPANs are not known to receive synaptic input and in fact have been characterized electrophysiologically by their lack thereof (*36*), some work has in fact suggested that AH-AH neuron interconnected pairs may exist (*37*). Immunohistochemical and electron microscopy investigation of synapses on enteric neurons further showed that calbindin-positive neurons, presumed IPANs, do receive synapses, though fewer than non-calbindin neurons, and some of those synapses were also calbindin-reactive (*38*). These observations informed a proposed “IPAN driver circuit” theory, in which IPANs form an interconnected network of positive feedback to synchronize and amplify sensory signaling and thus activate large swaths of enteric circuitry (*39*). In contrast, more recently, activation of large regions of the ENS has been suggested to be driven by interneuronal networks (*40*). It is important to note that our study was unsuccessful in localizing Cdh6 protein via immunohistochemistry, to confirm protein expression in neurons or in Cdh6-cre-tdT+ muscle cells or visualize synaptic connections between neurons or connections to other cell types. Further investigations are necessary to determine whether synaptic adhesion molecules, such as *Cdh6* and *Cdh8*, may in fact support IPAN-IPAN synapses underlying an “IPAN driver circuit.”

## Acknowledgments

We thank the members of the Kaltschmidt laboratory for experimental advice and discussions, Vanda Lennon (Mayo Clinic) for the HuC/D primary antibody and Beatriz G. Robinson for 3D printing of organ chambers for electrophysiology.

## Funding

National Institute of General Medical Sciences of the National Institutes of Health Award T32GM120007 (JGF)

National Institutes of Health Grant R01 EY030138 (XD)

National Health and Medical Research Council (NHMRC) project grant 1156416 (NJS)

Australian Research Council (ARC) Discovery Project grant DP220100070 (NJS)

NINDS 5RO1NS34774 (JRH)

Wu Tsai Neurosciences Institute, Stanford University (JAK)

Department of Neurosurgery, Stanford University (JAK)

National Institutes of Health Grant R21 HD110950 (JAK)

The Firmenich Foundation (JAK)

The Carol and Eugene Ludwig Family Foundation (JAK)

Stanford ADRC Developmental Project Grant (National Institutes of Health Grant P30AG066515) (JAK)

## Author contributions

Conceptualization: JGF, JAK

Resources: NJS, JRH, JAK

Methodology: JGF, GD, XD, TJH

Investigation: JGF, GD, LT, MAK, TJH

Visualization: JGF, GD, TJH

Funding acquisition: NJS, JRH, JAK

Supervision: NJS, JRH, JAK

Writing – original draft: JGF, GD

Writing – review & editing: JGF, NJS, JRH, JAK

## Data and materials availability

The data that support the findings of this study are available from the corresponding author upon reasonable request.

## Materials and Methods

### Mice

All procedures conformed to the National Institutes of Health Guidelines for the Care and Use of Laboratory Animals and were approved by the Stanford University Administrative Panel on Laboratory Animal Care. Mice were group housed up to a maximum of five adults per cage. Food and water were provided ad libitum and mice were maintained on a 12:12 LD cycle. Male and female mice were used in all experiments.

Wild-type C57BL/6J mice (#000664), Hb9:GFP mice (#005029), Ai14 (#007908), and Ai32 (#024109) (*21*) mice were obtained from the Jackson Laboratory and from the Animal Resource Center (ARC) in Western Australia, with JAX heritage. Cdh6^CreER^ mice (#029428) (*20*) were provided by Xin Duan (UCSF). Cdh6^CreER^ mice were crossed to Ai14 mice and Ai32 mice to generate mice heterozygous for each allele, termed Cdh6^CreER^;Ai14 and Cdh6^CreER^;Ai32. Tamoxifen (20 mg/mL in corn oil) was administered via oral gavage to a final dose of 2.5mg/10g mouse for five consecutive days beginning at 5-8 weeks of age. Induced mice were group housed for at least four weeks prior to experiments.

Adult male and female mice (Cdh6^CreER+^;Ai32 and Cdh6^CreER-^;Ai32) aged 16 to 19 weeks were euthanised by isoflurane inhalation overdose in accordance with Flinders Animal Welfare Committee guidelines (ethics approval #4004). The protocol for animal euthanasia is approved by the National Health and Medical Research Council (NHMRC) Australian code for the care and use of animal for scientific purposes (8^th^edition, 2013) and recommendations from the NHMRC Guidelines to promote the wellbeing of animals used for scientific purposes (2008).

### Dissection

Mice were culled by CO_2_ and cervical dislocation. Small intestine and colon were removed and flushed with ice cold PBS, then placed in a Sylgard-lined Petri dish with ice cold PBS for further dissection.

#### Wholemount preparations

Intestinal segments were prepared and fixed as in Gomez-Frittelli *et al* (*41*). Briefly, each intestinal segment was opened along the mesentery border, pinned flat under light tension serosa-side up, and fixed in 4% PFA in PBS at 4°C with gentle rocking for 90 minutes. Segments were washed three times in PBS at 4°C for at least 10 minutes with gentle rocking. Muscularis was separated from the mucosa at one end of the segment with fine forceps for 2-3 mm, then pinned mucosa-side up in the dish. The mucosa was peeled away from the muscularis with fine forceps while the muscularis was gently held down in the dish with a cotton swab. For immunohistochemistry, segments were processed immediately or stored in PBS with 0.1% NaN_3_ at 4°C until use. For RNAscope, muscularis segments were post-fixed in 4% PFA in PBS at 4°C with gentle rocking overnight, then washed three times in PBS at 4°C with gentle rocking for at least 10 minutes each wash before use.

### Immunohistochemistry

Immunohistochemistry was performed as described previously (*42*). Briefly, muscularis wholemount tissue segments about 7mm x 7mm were incubated with primary antibodies in PBT (PBS with 1% BSA and 0.3% Triton-X 100) at 4°C with gentle rocking overnight, then washed three times in PBT for at least ten minutes each at room temperature with gentle shaking. Tissues were incubated in secondary antibodies in PBT for 2 hours with gentle shaking, washed twice in PBT, and twice in PBS, then mounted on Superfrost Plus slides with Fluoromount G medium (Southern Biotech). Primary antibodies: human anti-HuC/D (1:75k) (gift from Vanda Lennon); sheep anti-GFP (1:1k) (Biogenesis); rabbit anti-RFP (1:1k) (Rockland); rabbit anti-PGP9.5 (1:4k) (Abcam). Secondary antibodies: donkey anti-human AlexaFluor (AF)-647 (1:500); donkey anti-sheep AF-488 (1:1k); donkey anti-rabbit AF-488 (1:1k); streptavidin AF-546 (1:500).

### RNAscope

*In situ* hybridization in combination with IHC was performed on muscularis wholemount tissues using the RNAscope Multiplex Fluorescent V2 Assay kit with RNA-Protein Co-detection Ancillary Kit [ACD], according to the manufacturer’s instructions with modifications as previously described (*43*). Probes used were *Cdh6* (#519541), *Cdh8* (#485461), *Nmu* (#446831), *Calcb* (#425511), and *eGFP* (#400281).

### Confocal imaging

Images were acquired on a Leica SP8 confocal microscope using a 20x (NA 0.75) oil objective at 1024 x 1024 pixel resolution. Tiled images (24-30 tiles) of Z-stacks (2.5µm between planes) were acquired and stitched together using the Navigator mode within LASX (Leica). Imaged regions were located away from the mesenteric border.

### Image analysis and quantification

Image analysis was performed using ImageJ/FIJI (NIH, Bethesda, MD), as described previously (*42*). HuC/D images (z-stack individual planes) were blurred and thresholded; then maximally projected and total neurons counted using the Analyze Particles function. Cdh6^CreER^;Ai32 expression and RNAscope *in situ* hybridization and Cdh6^CreER^;Ai14 expression were counted manually. Cell tracing was performed in Imaris using Filament Tracer (Bitplane, Oxford Instruments).

### Electrophysiological recordings

Whole cell patch clamp electrophysiological recordings of Hb9:GFP+ neurons were performed according to Osorio & Delmas (*14*), with modifications for recording from the distal colon. The protocol is described in brief below.

#### Tissue dissection and preparation

Mice aged 8-10 weeks were culled by CO_2_ and cervical dislocation. The colon was removed and flushed with ice cold oxygenated Krebs solution [118 mM NaCl, 4.8 mM KCl, 1 mM NaH2PO4, 25 mM NaHCO3, 1.2 mM MgCl2, 2.5 mM CaCl2 and 11 mM glucose, supplemented with scopolamine (2 M) and nicardipine (6 μM)], then placed in a Sylgard-lined Petri dish with ice cold oxygenated Krebs solution for further dissection. Krebs solution was changed out for fresh oxygenated solution every 5 minutes. Under a dissection microscope, the distal colon was pinned and the mucosa peeled away using fine forceps, leaving a few millimeters of mucosa along the edges of the tissue for pinning stability. The muscularis was then flipped over and re-pinned, serosa side up, and the longitudinal muscle carefully peeled away. The tissue was transferred to a custom 3D-printed recording chamber lined with a thin layer of clear Sylgard, and re-pinned under light tension, with the myenteric plexus facing up. The tissue was kept at 32°C and was continuously perfused with oxygenated Krebs solution. Hb9:GFP+ neurons were visually identified within a ganglion under epifluorescence illumination with a 455nM LED (Thorlabs, M455L2) and a 470 (excitation)/525 (emission) nm wavelength filter set. A local perfusion of protease XIV (0.2% in Krebs) (Sigma, P5417) was applied on top of the targeted cell to digest any muscle fiber residue. A 1-2MΩ pipet with a trimmed arm hair glued to the tip was used to brush and clean the surface of the ganglion. Further cleaning with 1mg/mL collagenase (Worthington, CLS-4) 4mg/mL dispase (Sigma, D4693) in Krebs solution was also performed to expose the GFP neuron for patching.

#### Patching and recording

Patch pipettes (4–6 MΩ) pulled from borosilicate glass were filled with internal solution containing in mM: 144 K-gluconate, 3 MgCl2, 0.5 EGTA, 10 HEPES, pH 7.2 (285/295 mOsm) and 2% biocytin (Millipore Sigma, B4261-100MG). Patch clamp recordings were collected with a Multiclamp 700A (Molecular Devices) amplifier, a Digidata 1440 digitizer and pClamp10.7 (Molecular Devices). Recordings were sampled and filtered at 10kHz. Passive properties analysis was performed using pClamp10.7. Analysis of action potentials was performed using a custom MATLAB (MathWorks) software. All recordings were performed at 32°C. Membrane potentials were not corrected for liquid junction potential. Immediately after whole cell configuration, the cell was maintained at −70mV and a short voltage clamp membrane test protocol consisting of 20 times 600ms, 10mV depolarization steps was performed to assess cell health and recording conditions. Recordings were performed in Hb9:GFP+ colonic neurons with an access resistance less than 30MΩ (16.69 ± 2.63 MΩ). Next, the current clamp mode was used to measure resting membrane potential (RMP), input resistance (Rin) and APs stimulated. Membrane potential was not adjusted from resting potential, and cells were depolarized by 1 second current pulses in 10pA increments until APs were triggered (rheobase). Finally, if the seal was still stable, a voltage clamp steady-state inactivation of T-current protocol was performed as previously described (*44*). In brief, a sequence of depolarization from −90 to −45mV for 500ms quickly followed by a depolarization to −40mV for 200ms. Tissues were then fixed and immunostained according to *Wholemount preparations* and <u>Immunohistochemistry</u> sections above.

### Mechanical recordings & optical stimulation

Optogenetic stimulation experiments were performed as previously described (*29*). A 2.5 mm stainless steel rod was inserted through the lumen of the colon and mounted in an organ bath (120 * 40 * 12 mm; L*W*H) located on a heated base. Krebs solution (35.5-36°C) superfused the bath (∼5 mL/min). Smooth muscle force was recorded via 4 evenly spaced hooks in the colonic muscularis externa, each linked to an isometric force transducer (Grass FT03C) by suture thread. Initial base resting tension was set between 0.5 - 1.0 g. Preamplified signals (Biomedical Engineering, Flinders University) were digitized by a PowerLab 16/35 (ADInstruments, Bella Vista, NSW, Australia) and recorded using LabChart 7 software (ADInstruments) on iMac computer. Post hoc analysis of the mechanical recordings was done using LabChart 8 software on PC.

For optical stimulation during mechanical recordings *in vitro*, two LEDs (emitting 470 nmλ photons; C470DA2432, Cree Inc., NC, USA) were used, driven by a variable power supply. The area of light emission from each LED was 240 μm x 320 μm (0.0768 mm^2^). To characterise LED function, light power density across a range of currents was measured 5 mm from the LED using a standard photodiode power sensor (S120C; Thorlabs, NJ, USA) and a power meter (Thorlabs, PM100USB). The stimulator panel within LabChart software was used to set parameters and manually trigger LED pulse trains via the 10V analogue output of the PowerLab and an ILD1 opto-isolator.

### Intraluminal pellet CMC recordings

To record proximal and distal colon CMCs separately (*45*, *46*), full length colon was bisected halfway between the caeco-colonic junction and terminal rectum, creating equal length proximal and distal colon preparations. Each preparation was suspended vertically on a stainless-steel holder inside a glass, water jacketed organ bath containing Krebs solution (Fig.6J, K). A 2.7 mm diameter synthetic pellet (polymethyl methacrylate, “Perspex”) was placed inside the gut lumen and linked by stainless steel rod to a force transducer (MLT0420, ADInstruments), allowing measurement of both anterograde and retrograde propulsive forces on the pellet. Signals were amplified by bridge amplifier (FE224, ADInstruments), digitized at 1kHz (PLCF1, ADInstruments) and recorded using LabChart 8 software.

ZD7288 (73777, Sigma-Aldrich) was dissolved in water as stock solution at 10 mM. Caesium Chloride (C4036, Sigma-Aldrich) was dissolved in water as stock solution at 200 mM. Tetrodotoxin citrate (T-550, Alomone Labs) was dissolved in water as stock solution at 3 mM. Control, ZD7288, CsCl and washout periods were at least 30 mins; TTX was applied for at least 10 minutes.

### Statistical analysis

Statistical tests and graphical representation of data were performed using Prism 9 software (GraphPad). Statistical comparisons were performed using paired t tests (one-tailed, CMC intervals; two-tailed, peak amplitude, AUC, duration) and Welch’s t test (marker colocalizations). Asterisks indicate significant differences.

---

**Fig. S1.**
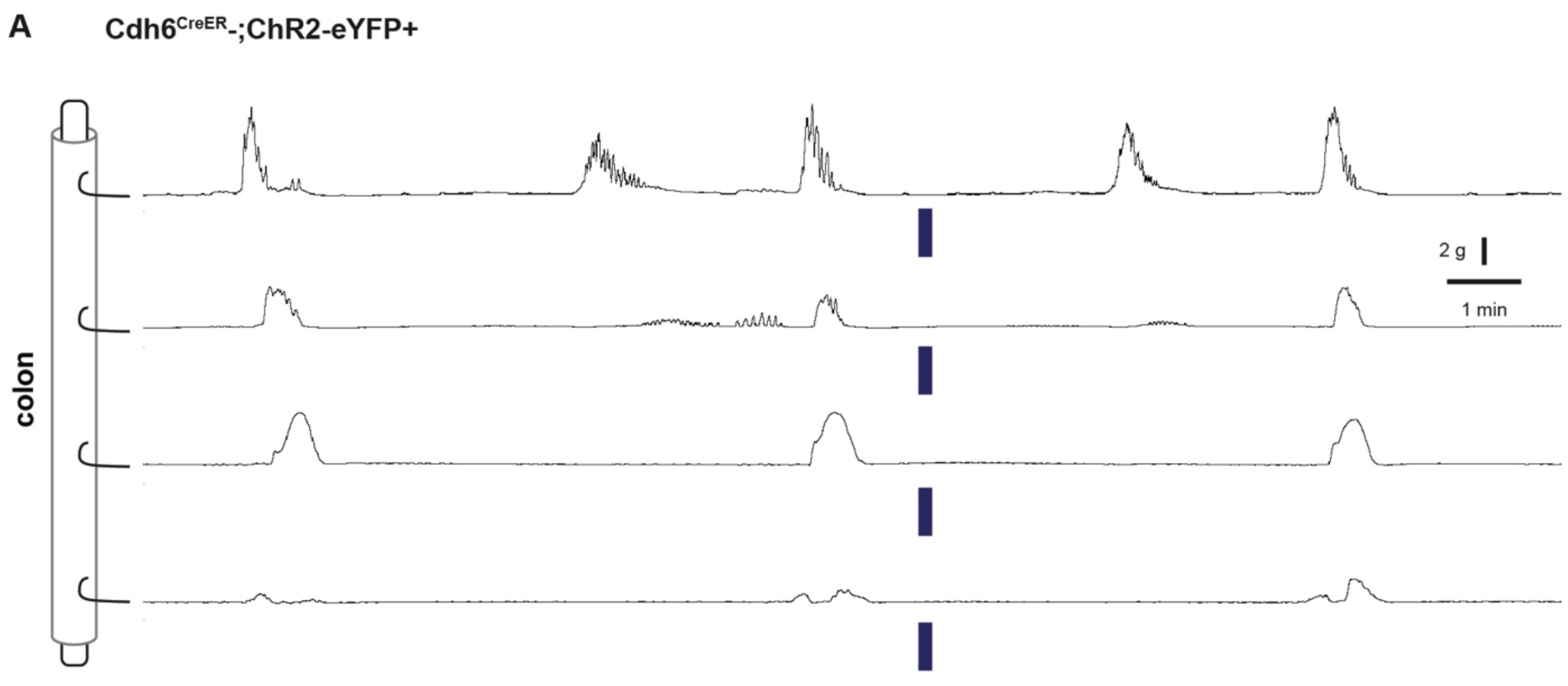
Optogenetic stimulation control. (**A**) Representative force traces of control Cdh6^CreER^-;Chr2eYFP+ colon (n = 5). Blue bars indicate timing of light stimulation. LEDs placed distal to distal hook.

**Fig. S2.**
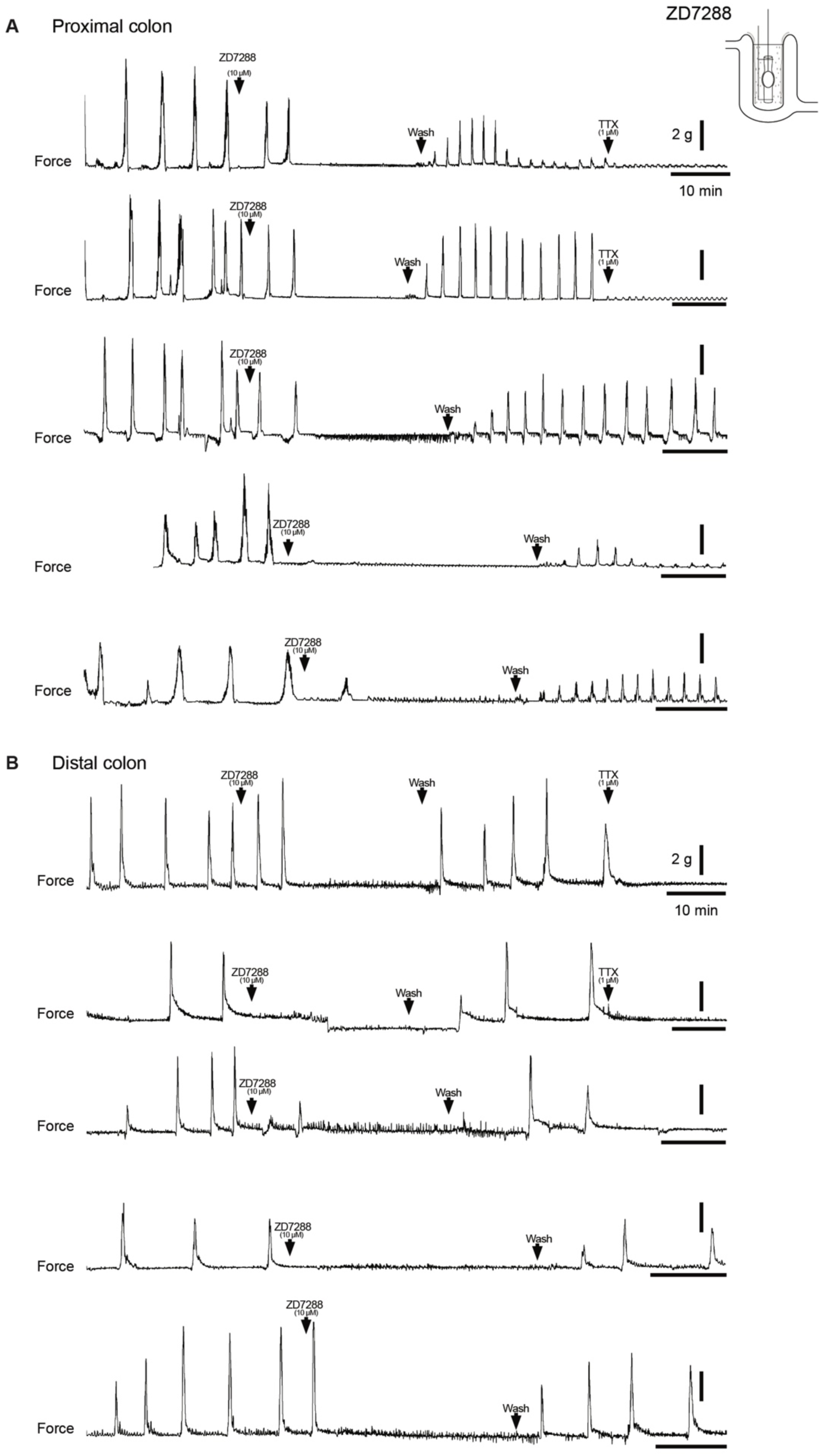
Pharmacologic blockade of *I_H_* with ZD7288 abolishes spontaneous CMCs. (**A, B**) Representative force traces from tethered pellet in proximal half (A) or distal half (B) of colon. Addition of 10 µM ZD7288 (first arrowhead), followed by washout in Krebs (second arrowhead), and addition of 1 µM TTX (third arrowhead). Scale bars represent 2 g force (vertical bars) and 10 minutes (horizontal bars) for all traces.

**Fig. S3.**
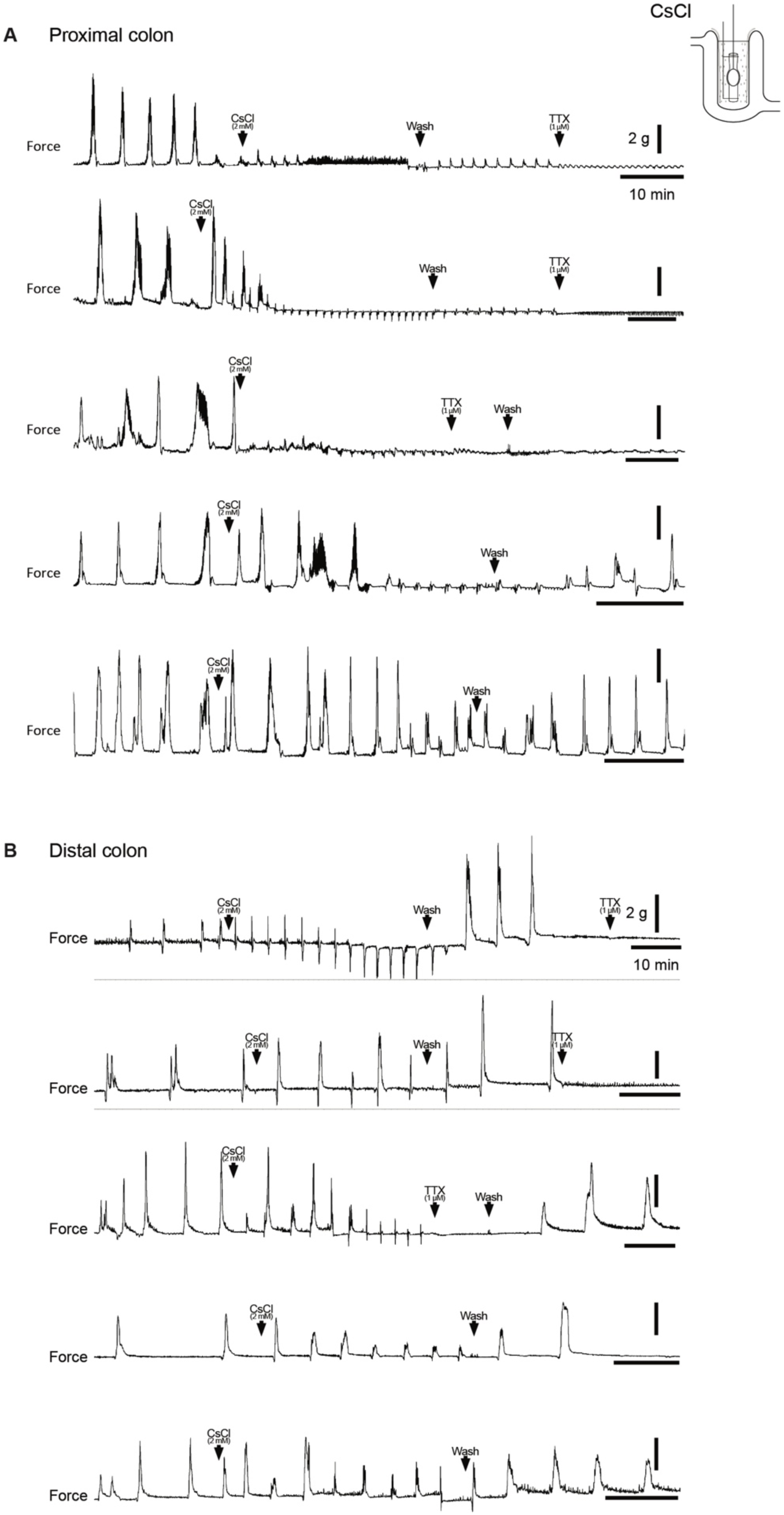
Pharmacologic blockade of *I_H_* with CsCl impairs generation of CMCs. **(A, B)** Representative force traces from tethered pellet in proximal half (A) or distal half (B) of colon. Addition of 2 mM CsCl (first arrowhead), followed by washout in Krebs (second arrowhead), and addition of 1 µM TTX (third arrowhead). Scale bars represent 2 g force (vertical bars) and 10 minutes (horizontal bars) for all traces.

